# A Joint Model of RNA Expression and Surface Protein Abundance in Single Cells

**DOI:** 10.1101/791947

**Authors:** Adam Gayoso, Romain Lopez, Zoë Steier, Jeffrey Regier, Aaron Streets, Nir Yosef

**Affiliations:** Center for Computational Biology, University of California, Berkeley; Department of Electrical Engineering and Computer Sciences, University of California, Berkeley; Department of Bioengineering, University of California, Berkeley; Department of Statistics, University of Michigan, Ann Arbor; Chan Zuckerberg Biohub, San Francisco, California; Ragon Institute of MGH, MIT and Harvard

## Abstract

Cellular indexing of transcriptomes and epitopes by sequencing (CITE-seq) combines unbiased single-cell transcriptome measurements with surface protein quantification comparable to flow cytometry, the gold standard for cell type identification. However, current analysis pipelines cannot address the two primary challenges of CITE-seq data: combining both modalities in a shared latent space that harnesses the power of the paired measurements, and handling the technical artifacts of the protein measurement, which is obscured by non-negligible background noise. Here we present Total Variational Inference (totalVI), a fully probabilistic end-to-end framework for normalizing and analyzing CITE-seq data, based on a hierarchical Bayesian model. In totalVI, the mRNA and protein measurements for each cell are generated from a low-dimensional latent random variable unique to that cell, representing its cellular state. totalVI uses deep neural networks to specify conditional distributions. By leveraging advances in stochastic variational inference, it scales easily to millions of cells. Explicit modeling of nuisance factors enables totalVI to produce denoised data in both domains, as well as a batch-corrected latent representation of cells for downstream analysis tasks.

## 1 Introduction

Single-cell RNA sequencing (scRNA-seq) enables comprehensive quantitative characterization of a cell’s mRNA profile and has resulted in novel findings about the molecular circuitry of cell populations [1, 2]. Extending scRNA-seq, cellular indexing of transcriptomes and epitopes by sequencing (CITE-seq) simultaneously measures the abundance of selected proteins on the cell surface with results comparable to gold-standard flow cytometry [3]. As surface proteins are routinely used as a measure of cell phenotype, CITE-seq provides an exciting new opportunity for enhancing the quality of data interpretation [3, 4, 5], especially as the number of assayed proteins per experiment grows (with over 300 barcoded-antibodies commercially available [6]). Recent approaches to CITE-seq data analysis consist of deriving clusters based on the mRNA data and mapping protein values to these clusters for the task of cell-type labeling. However, this strategy assumes that cell-cell similarities depend only on mRNA, and neglects the distinctive information contained in the protein measurements. We aim to combine both measurements into one representation of cell state, while addressing the unique technical biases of each modality.

A reasonable modeling assumption is that both mRNA and protein counts are generated from a low-dimensional manifold of cellular states [1]. While a flourishing body of research has focused on using generative models to learn a biologically meaningful low-dimensional representation of cells in scRNA-seq datasets [7, 8, 9, 10], no proposed method can additionally handle the complexities of the protein counts. CITE-seq protein counts are overdispersed like mRNA counts, but do not suffer from limited capture efficiency. Instead, the protein counts are obscured by a non-negligible background signal, which may arise due to non-specific binding of antibody probes and/or ambient antibodies. As a result, the distribution of protein counts across cells is often bimodal with a background and foreground component. A natural way to preprocess the protein counts is to fit a mixture model to each protein globally and replace a count with its probability of being generated from the larger component [11, 12]. However, there is no basis for assuming global bimodality of protein counts since a dataset typically comprises heterogeneous populations of cells each with distinct surface proteomes. Consequently, using the same signal-noise decision boundary for all cells can be inappropriate.

Here we propose Total Variational Inference (totalVI), a coupled generative model and inference procedure for CITE-seq data, which addresses these issues. In totalVI, both mRNA and protein counts of a cell are assumed to be random variables generated from a low-dimensional latent variable that represents the underlying biological state of a cell and contains information from both domains. Such a framework enables end-to-end analysis of this data – a joint batch-corrected latent representation (for stratifying cells into types), denoised data in both domains, and differential expression of genes and proteins. totalVI leverages advances in stochastic optimization and easily scales to millions of cells.

## 2 The totalVI probabilistic model

A CITE-seq experiment produces two vectors for a cell *n, x*_*n*_ and *y*_*n*_, where *x*_*nr*_ is the number of mRNA molecules detected for gene *r* and *y*_*nt*_ is the number of cell surface molecules detected for protein *t*. Furthermore, a dataset has *R* genes and *T* assayed proteins. Let *s*_*n*_ be the batch cell *n* was processed in (one-hot encoded) with a total of *B* batches.

Let *z*_*n*_ be a latent variable describing the biological state of cell *n*. Given *z*_*n*_ and a cell-specific scaling factor *𝓁*_*n*_ representing technical factors like the mRNA sequencing depth, *x*_*nr*_ follows a negative binomial distribution with gene-specific i nversed ispersion and w ith t he prior for *𝓁*_*n*_ set as in [7]. Let *µ*_*nt*_ be a latent variable representing the mean of the background distribution for the protein counts, sampled from a protein-batch-specific prior. Given *z*_*n*_ and *µ*_*nt*_, we model *y*_*nt*_ as a negative binomial mixture to capture observed protein counts arising from the background or foreground. The mean parameters of the mixture components are structured such that the foreground mean is strictly larger than the background mean, which also identifies the mixture as the inverse dispersion parameter is shared between components.

The full generative process is outlined in Algorithm 1. The prior parameters for *µ*_*nt*_ are learned in a variational Bayesian inference fashion. The neural networks are dense with one hidden layer, ReLU activations, batch normalization and dropout. Notably, *z*_*n*_ follows a logistic normal distribution, meaning cells can be interpreted as having “membership” to dimensions of the latent space and that archetypal analysis can be performed (not shown) [13, 14].

## 3 Inference

We use variational inference to obtain the approximate posterior distribution

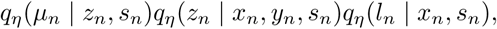

where *η* is the set of parameters of an inference network – a neural network that takes a cell’s combined expression as input and outputs the parameters of the approximate posterior. The variational distribution *q*_*η*_(*µ*_*n*_ | *z*_*n*_) has parameters specific to the cell *n* and not global parameters like the prior. We optimize the evidence lower bound (ELBO) [15] of log *p*_*ν*_ (*x*_1:*N*_, *y*_1:*N*_ | *s*_1:*N*_) with respect to the variational parameters *η* and model parameters ν using stochastic gradients [16]. To avoid inference over discrete random variables, we analytically integrate out *v*_*nt*_, yielding *p*_*ν*_ (*y*_*nt*_ | *z*_*n*_, *µ*_*nt*_), which is a mixture of negative binomials. We use the Adam optimizer [17], along with deterministic warm-up, and a reduction of the learning rate upon plateau of the ELBO on a validation set.

### Algorithm 1

The totalVI generative model. The negative binomial distribution is parameterized by its mean and inverse dispersion. Let *ν* be the set of model parameters described here.

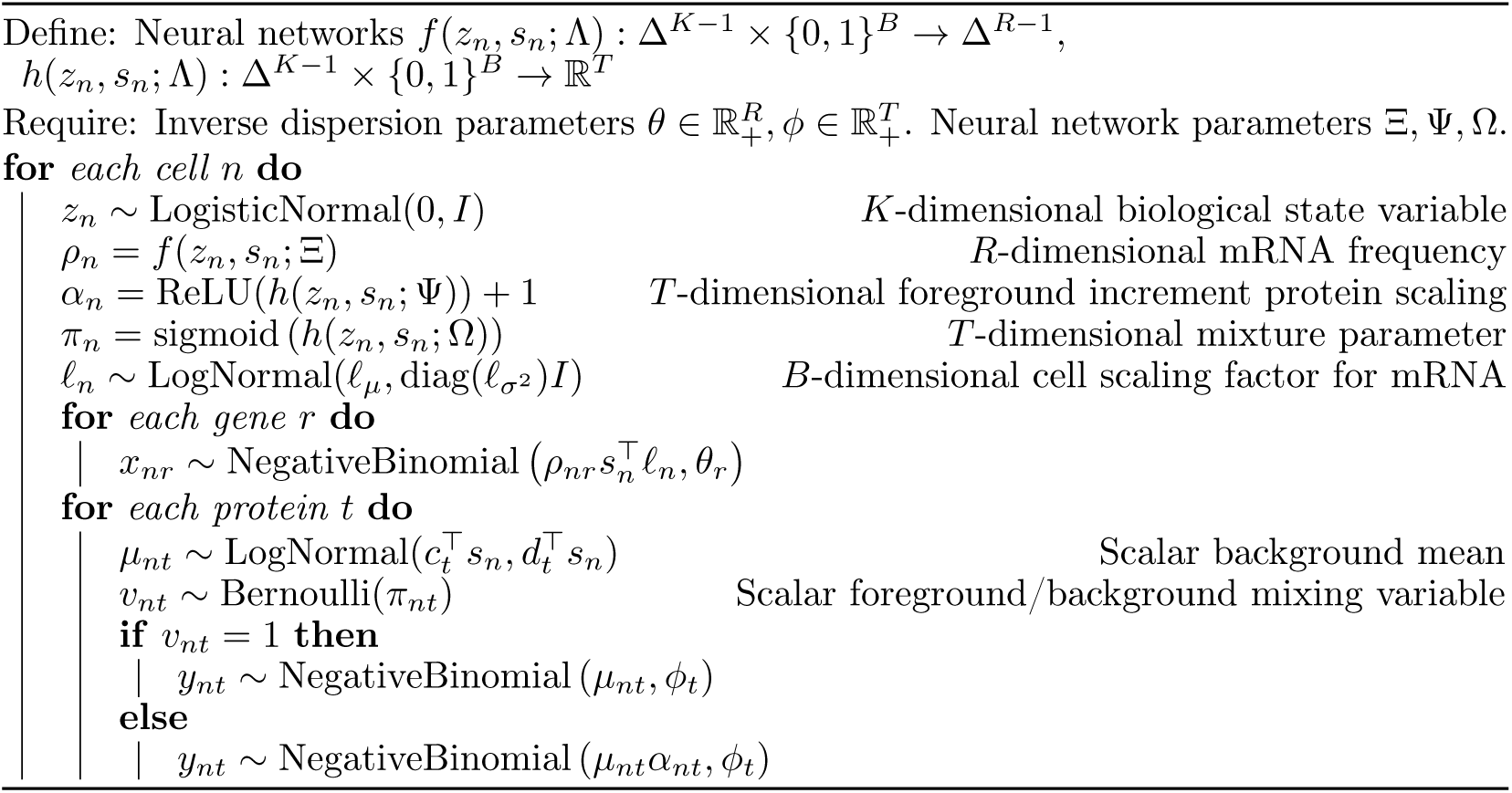

## 4 Performance benchmarks

We assess the performance of totalVI on two tasks: generalization to held-out data and posterior predictive checks (PPC) of coefficient of variation. We compare totalVI to factor analysis (FA) in which we either fit on the data that is logarithmized plus one (log), or log normalized data where the two modalities are independently normalized by their library size prior to log transformation (log rate). We also compare to the state-of-the-art scRNA-seq model, scVI [7], where we treat the protein data as additional features (genes) and use a negative binomial likelihood distribution for both mRNA and protein counts.

For each of these tasks, we use two datasets: (1) 7,225 peripheral blood mononuclear cells (PBMC10k) from 10X Genomics [18] and (2) 8,412 cells from a MALT tumor (MALT) [19]. We first remove genes that are expressed in fewer than 1% of cells and retain the top 5,000 genes as measured by variance across cells. Both datasets contain 14 proteins. We filter cells that were below the first percentile for protein UMI counts and had expressed fewer than 500 genes. For PBMC10k we filter doublets using DoubletDetection [20].

### Generalization to held-out data

We compare totalVI to scVI and the FA baselines using 20 latent dimensions for each method. For each model, we compute separately for protein and gene features the mean squared logarithmic error (MSLE) between the observed values and the mean of the posterior predictive distribution on a held-out subset representing 6% of the total dataset. For example,

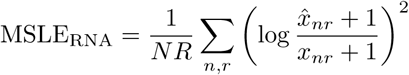

where 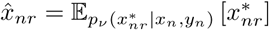, the mean of the posterior predictive, while MLSE_Protein_ is computed using *y*_1:*N*_ instead. Table 1 shows that totalVI has lower MSLE with respect to mRNA and proteins, which can be attributed to the superior noise model for proteins used in totalVI.

**Table 1:** Mean squared logarithmic error between observations and the mean of the posterior predictive distribution.

| Model | PBMC10k |  | MALT |  |
| --- | --- | --- | --- | --- |
|  | Protein | RNA | Protein | RNA |
| FA (log) | 0.935 | 0.167 | 0.938 | 0.169 |
| FA (log rate) | 0.860 | 0.125 | 0.870 | 0.121 |
| scVI | 0.739 | 0.105 | 0.474 | 0.103 |
| totalVI | **0.599** | **0.103** | **0.431** | **0.098** |

We also compute the held-out log-likelihood for totalVI and scVI. Table 2 shows that totalVI outperforms scVI on both datasets. As log-likelihoods for models with discrete and continuous likelihoods are not directly comparable, we computed the calibration error [21] in order to quantify the quality of each model’s uncertainty estimates. totalVI and scVI have lower calibration error than FA models (results not shown).

**Table 2:** Negative log likelihood on held-out data.

| Model | PBMC10k | MALT |
| --- | --- | --- |
| scVI | 3349.44 | 3180.80 |
| totalVI | **3337.89** | **3158.78** |

### Posterior predictive checks

We perform a PPC [22] of the coefficient of variation (CV) for each protein and each gene. For each model, we sample the posterior predictive distribution 25 times, calculating the CV for each sample and for each feature. After averaging over samples, we separate the predicted CVs by feature type (mRNA or protein), resulting in two vectors: CV_RNA_ and CV_Protein_. We report the median absolute error between the observed and the predicted CV_RNA_ and CV_Protein_. Table 3 shows that totalVI outperforms other methods in both modalities indicating better model fit.

**Table 3:**
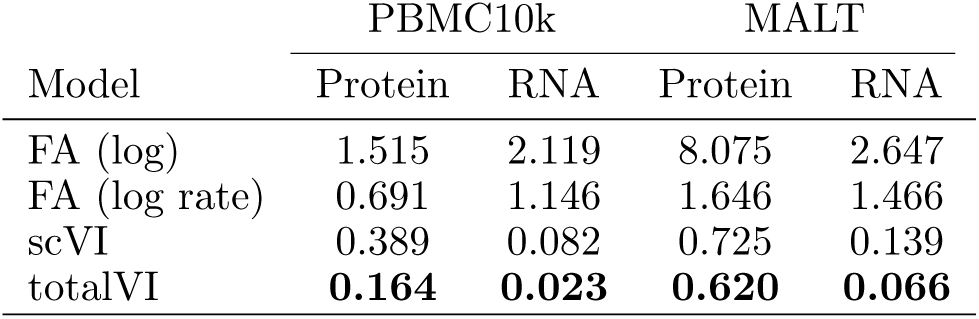
CV PPC. Median absolute error.

## 5 Protein background disentanglement

Disentangling background and foreground protein counts is crucial for mitigating spurious differential expression and cell-type labeling results. Previous work derives a linear cutoff on the number of counts for each protein based on spiked-in cells that do not express the proteins specifically recognized by the barcoded antibodies [3]. Others fit mixture models to each protein, which assumes that all cells are subject to the same background distribution [11, 12]. Our approach, which obviates the need for negative control cells or the assumption of a constant background distribution, models each *y*_*nt*_ | *z*_*n*_, *µ*_*nt*_ as a negative binomial mixture, where the Bernoulli parameter *π*_*nt*_ | *z*_*n*_ can be interpreted as the probability that a cell’s protein count came from the background. Thus, decision boundaries are cell-protein-specific, taking into account the overall state of the cell (genes and proteins).

As an example, consider the CD16 protein in PBMC10k (Figure 1). We highlight a subset of cells that could be called background by a global mixture model but are predicted to be foreground by totalVI (blue: 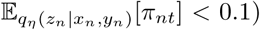. We also highlight a subset of cells with similar magnitude of expression as the blue set but with higher predicted background probability (red: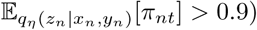). Consistent with the model, we observe that the foreground-predicted subset (blue) corresponds to natural killer cells and CD16+ monocytes (both of which are known to express CD16), while the background subset of cells (red) correspond to the other PBMC cell types (which do not tend to express CD16; note that cell-types were determined by mRNA).

**Figure 1:**
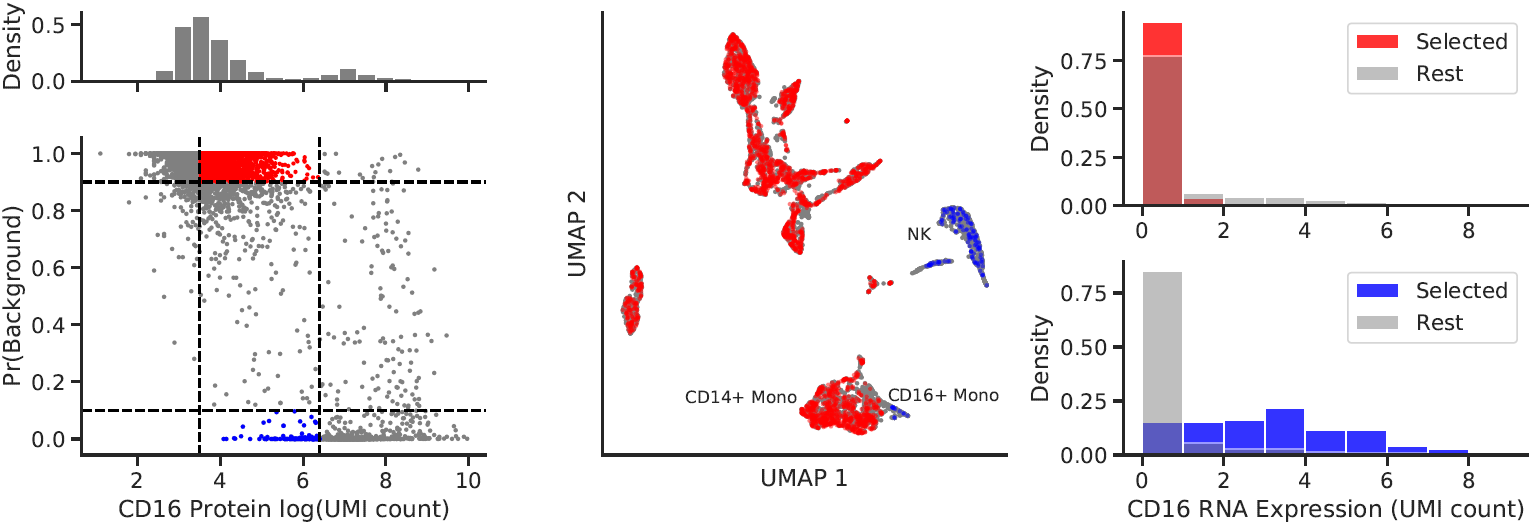
Investigation of totalVI background prediction for CD16 protein counts in PBMC10k.

Figure 1 also shows that mRNA counts of the CD16 gene are high in the foreground and low in the background subset. Thus, totalVI makes a non-trivial prediction that goes beyond a simple cutoff – by leveraging information between all cells, genes, and proteins.

Despite using a two-component mixture density, totalVI can also disentangle the background of proteins that are trimodal globally. For example, it has been shown using flow cytometry that monocytes have fewer CD4 protein molecules on their surface relative to CD4+ T-cells [23] and that other PBMC types do not tend to express CD4 on their surface. As such, the distribution of the CD4 protein in PBMC10k is trimodal (lowest mode corresponding to background).

Figure 2 shows the trimodal distribution of CD4 as well as the probability of background mapped to a UMAP [24] projection of 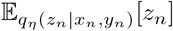. The cells we manually labeled (based on mRNA) as CD4+ T-cells and monocytes are indeed determined to have been mostly generated from the foreground component, while the remaining cells fall within the background part. This result is made possible due to the mixture being conditionally dependent on *z*_*n*_, thus defining the two modes of the foreground-background dichotomy in a manner local to the latent space. These predictions will be critical for quantifying differential protein expression.

**Figure 2:**
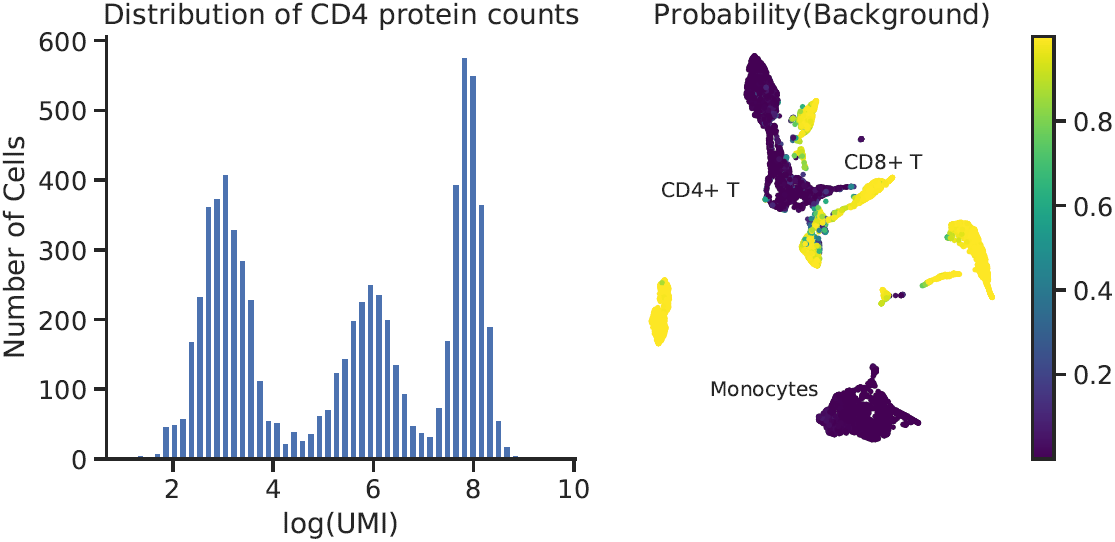
CD4 Protein disentanglement in PBMC10k. (Left) Distribution of log counts for CD4 protein. (Right) 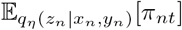 projected on UMAP.

## 6 Data denoising

Another application of totalVI is data denoising, in which a denoised expression matrix (mRNA and protein) is produced that can then be used as input for other downstream tasks, like building a feature-wise correlation matrix to identify signaling and regulatory networks, or examining the dynamics of transcription and translation. The totalVI denoised expression matrix is constructed by first replacing *x*_*nr*_ with 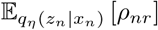 for all cells and all genes. For a protein count *y*_*nt*_, we replace with the quantity 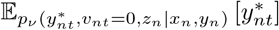 which is interpreted as the expected protein count given it was generated by the foreground component of the mixture, adjusted for the probability it was generated by the foreground. It is then normalized so that the denoised protein expression for a cell *n* resides in the simplex. Thus, protein counts likely to have been generated from the background will have magnitude near zero in the denoised matrix.

We construct Spearman correlation matrices between all proteins and 500 genes chosen randomly except for the inclusion of genes that encode for the assayed proteins. The matrices are derived using (1) denoised expression, (2) raw counts, (3) posterior predictive counts.

In Figure 3 we plot the posterior predictive correlations relative to the raw correlations as well as the denoised correlations relative to the raw correlations. Correlations of features derived from the same gene are in red while all others are in grey. We emphasize that totalVI has no prior knowledge of RNA-protein translation relationships. We see that the denoised correlations are more extreme than their raw counterparts, however posterior predictive correlations largely match the original raw correlations, indicating that extreme denoised correlations are not symptomatic of fitting a low-dimensional model and that totalVI is instead likely to restore biological correlations.

**Figure 3:**
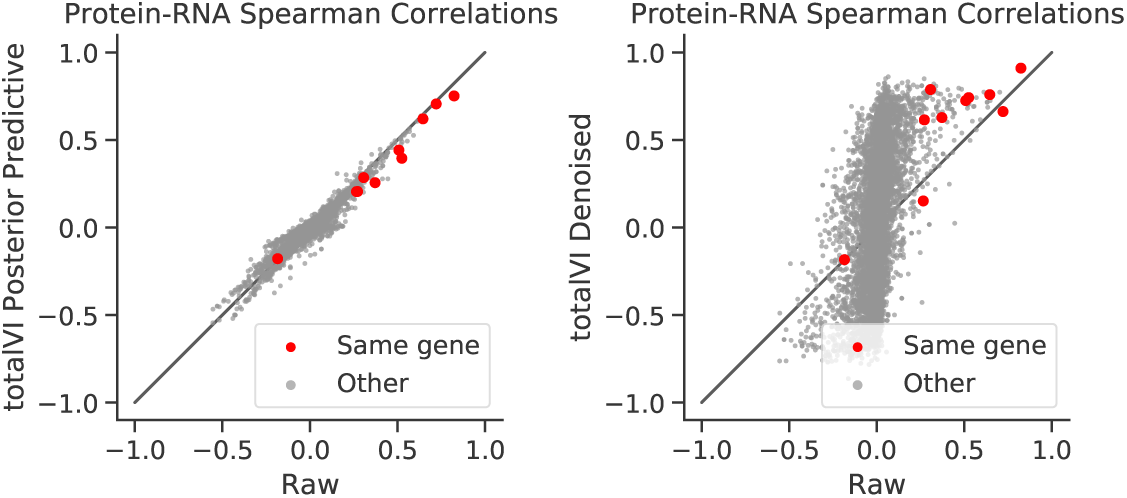
totalVI posterior predictive and denoised Spearman correlations versus raw correlations in PBMC10k.

## 7 Dataset harmonization

We demonstrate how totalVI can be used to generate a batch-corrected joint latent representation by harmonizing two PBMC datasets (PBMC10k and another dataset of 5k PBMCs from 10X Genomics [25] with genes and proteins subsetted to match PBMC10k; results in Figure 4). totalVI has a unique advantage over popular methods based on mutual nearest neighbors [4, 26], as no similarity metric between cells is necessary, which may be biased toward one modality. Instead, independence between *z*_*n*_ and *s*_*n*_ is a byproduct of the invariance of the prior on *z*_*n*_ to the batch. Another benefit of our method is the ability to marginalize over batch in order to generate batch-free denoised values, or perform differential expression over batches.

**Figure 4:**
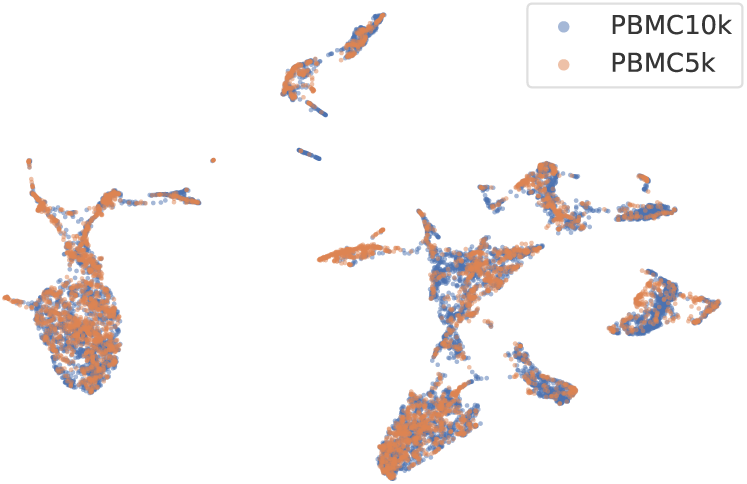
totalVI batch-corrected joint latent representation visualized with UMAP.

## Code availability

The implementation to reproduce the experiments of this paper is available at https://github.com/adamgayoso/totalVI_reproducibility. The reference implementation of totalVI is available at https://github.com/YosefLab/scVI. Datasets are publicly available from 10X Genomics.

## Acknowledgements

AG is supported by NIH Training Grant 5T32HG000047-19. ZS is supported by the NSF GRFP. This work is supported in part by NIH Grant R35GM124916. AS and NY are Chan Zuckerberg Biohub investigators.

